# Astrocytes modulate baroreceptor reflex sensitivity at the level of the nucleus of the solitary tract

**DOI:** 10.1101/725770

**Authors:** Svetlana Mastitskaya, Egor Turovsky, Nephtali Marina, Shefeeq M. Theparambil, Anna Hadjihambi, Andrew G. Ramage, Alexander V. Gourine, Patrick S. Hosford

## Abstract

Astrocytes play an important role in cardiovascular reflex integration at the level of the nucleus tractus solitarii (NTS). Existing reports from brainstem slice preparations suggest that astrocytes here respond to input from the solitary tract by increasing intracellular calcium. However, the physiological significance of this neuron-astrocyte signaling *in vivo* remains unknown. Here, we report that stimulation of the vagus nerve in an anesthetized rat induced rapid [Ca^2+^]_i_ increases in astrocytes transduced to express calcium sensor GCaMP6. The receptors involved were determined using brainstem-derived astroglial cell cultures were loaded with [Ca^2+^] indicator Fura-2. 5-HT (10 µM) caused robust increases in [Ca^2+^]_i_, and pharmacological interrogation revealed the expression of functional 5-HT_2A_ receptors. This observation was confirmed *in vivo*: intravenous administration of ketanserin decreased the magnitude of [Ca^2+^]_i_ responses, induced by vagal afferent simulation, by ∼50%. However, the response was completely blocked by topical application of the AMPA receptor antagonist CNQX alone. To investigate the role of astrocyte-neuron communication, the vesicular release in the NTS astrocytes was blocked by virally driven expression of a dominant-negative SNARE protein *in vivo.* This increased baroreflex sensitivity in awake animals, which was also observed in anesthetized animals after topical application of the P2Y_1_ receptor antagonist MDS-2500 to the NTS. We hypothesize that NTS astrocytes respond to incoming afferent release of glutamate and this response is modulated by 5-HT originating from vagal afferents or other sources. ATP is then released, which acts on inhibitory interneurons via P2Y_1_ receptors and thus modulates the expression of cardiovascular reflexes.

**Significance statement:** Cardiorespiratory nuclei in the brainstem integrate cardiovascular sensory information to optimise tissue perfusion and blood gas concentrations. We describe experimental evidence that NTS astrocytes participate in setting the baroreflex sensitivity by release of ATP acting on P2Y_1_ receptors on inhibitory interneurons. Activation of astrocytes is partly under control of 5-HT co-released with glutamate from vagal afferents, which allows modulation of autonomic response to high frequency/duration of afferent stimulation by monitoring extra-synaptic 5-HT acting on glial 5-HT_2A_ receptors. This could represent a signaling pathway that is activated under pathological conditions and is responsible for baroreflex impairment in conditions that result in astrogliosis, for example from systemic inflammatory response or chronic hypoxia/hypercapnia.

## Introduction

All fundamental reflexes essential for the maintenance of cardiorespiratory homeostasis are orchestrated by the autonomic circuits located in the lower brainstem. Effective operation of these reflexes ensures autonomic balance and maintains cardiovascular physiology. Impaired operation of these reflexes (the baroreflex in particular) may contribute to the development of cardiovascular disease and serves as a robust predictor of all-cause mortality (La Rovere et al., 1998; La Rovere et al., 2001; McCrory et al., 2016). Brainstem autonomic circuits receive sensory information via afferent fibers running within the 9^th^ (glossopharyngeal) and 10^th^ (vagus) cranial nerves. These afferents terminate in the nucleus of the solitary tract (NTS), located in the dorsal aspect of the brainstem, and release glutamate as the principal transmitter at the first central synapse (Talman, 1997; Baude et al., 2009).

Glutamatergic transmission in the NTS is modulated by a complex variety of other transmitters (see Sevoz-Couche & Brouillard, 2017), with 5-hydroxytryptamine (serotonin, 5-HT) considered to play a key role in this important cardiovascular sensory nucleus (Ramage and Villalón, 2008; Hosford and Ramage, 2019). The transmitters and receptors involved in neuron-neuron communication necessary for reflex integration have been well studied. However, the role of astrocytes in this brain area are much less understood. This is despite a notable abundance of astrocytes within the NTS (Dallaporta et al., 2018) and significant evidence that astrocytes modulate the activities of other CNS circuits – for example, learning and memory (Han et al., 2012; Navarrete et al., 2012), control of sleep (Halassa et al., 2009) and regulation of breathing (Gourine et al., 2010; Sheikhbahaei et al., 2018).

Two important previous studies give an insight into the potential role of astrocytes in cardiovascular sensory afferent processing. Firstly, electrical stimulation of the solitary tract in a brainstem slice preparation activated the NTS astrocytes (shown by an increase in intracellular calcium), and this was reduced by AMPA receptor antagonist NBQX (McDougal et al., 2011). In a second study, a non-specific toxin saporin was used to ablate astrocytes, which resulted in impairment of baroreflex sensitivity and alteration of chemo- and von Bezold-Jarisch reflexes. This reflex dysfunction then led to blood pressure lability and, in some animals, resulted in sudden cardiac death (Lin et al., 2013). Together, these findings indicate that NTS astrocytes can respond to vagal inputs and may play an important role in cardiovascular reflex expression. However, it is still unknown if NTS astrocytes release gliotransmitters when activated and, if so, the identity of these transmitters. Moreover, ablation of astrocytes also removes the structural and metabolic support they provide and thus could mask the subtleties of their role in transmission and subsequent effect on physiological mechanisms controlled by NTS circuits.

In the present study we sought to address these questions by performing *in vivo* calcium imaging in NTS astrocytes expressing a genetically encoded Ca^2+^ sensor under control of the GFAP promoter. Next, as 5-HT is also known to be released from vagal afferent terminals to modulate glutamatergic transmission (Jeggo et al., 2005; Oskutyte et al., 2009; Hosford et al., 2015), the presence of functional 5-HT receptors on NTS glia was explored using calcium imaging in cultured astrocytes and *in vivo*. Previously, 5-HT receptors have been shown to be present on astrocytes in many brain areas (Sanden et al., 2000) but have not been positively identified in the brainstem. Finally, we investigated how these signaling mechanisms interact to shape cardiovascular reflexes by blocking Ca^2+^-dependent vesicular release in NTS astrocytes of conscious adult rats and monitoring baroreflex sensitivity.

The data obtained suggest that in response to incoming afferent information NTS astrocytes are activated by glutamate (acting at AMPA receptors) and 5-HT acting at 5-HT_2A_ receptors. This causes the release of ATP, which restricts the expression of baroreflex via activation of P2Y_1_ receptors, presumably on NTS inhibitory interneurons.

## Materials and Methods

The experiments were performed in Sprague Dawley rats in accordance with the European Commission Directive 2010/63/EU (European Convention for the Protection of Vertebrate Animals used for Experimental and Other Scientific Purposes) and the United Kingdom Home Office (Scientific Procedures) Act (1986) with project approval from the Institutional Animal Care and Use Committee of the University College London.

### In vivo gene transfer

Young male Sprague Dawley rats (100-120 g) were anesthetized with a mixture of ketamine (60 mg of kg^−1^, i.m.) and medetomidine (250 μg kg^−1^, i.m.) and placed in a stereotaxic frame. NTS astrocytes were targeted bilaterally to express either a genetically encoded Ca^2+^ sensitive indicator GCaMP6 (to record activity) or dominant negative SNARE protein (dnSNARE) (to block vesicular exocytosis; Sheikhbahaei et al., 2018). Stable GCaMP6 expression along the rostro-caudal extent of the NTS was achieved by placing two microinjections per side (0.25 μl each, 0.1 μl min^−1^) of an adeno-associated viral vector (AAV) designed to express GCaMP6 under the control of an enhanced glial fibrillary acidic protein (GFAP) promoter (AAV5.GfaABC1D.cytoGCaMP6f.SV40, titer 7×10^9^ viral genomes/mL; Penn Vector Core). To block vesicular release mechanisms in the NTS astrocytes, adenoviral vector (AVV) with the same promoter was used to drive the expression of dnSNARE (AVV-sGFAP-dnSNARE-EGFP, titer 7.7×10^9^ viral genomes/mL). Detailed validation of dnSNARE efficacy in blocking vesicular release mechanisms in astrocytes was reported previously (Sheikhbahaei et al., 2018). NTS astrocytes of control animals were targeted with AVVs to express calcium translocating channelrhodopsin variant (CatCh) fused with EGFP (AVV-sGFAP-CatCh-EGFP, titer 2.1×10^9^ viral genomes/mL). Anesthesia was reversed with atipamezole (1 mg kg^−1^). No complications were observed after the surgery and the animals gained weight normally.

### Anesthetized animal preparation and calcium imaging in NTS astrocytes in vivo

The experiments were conducted four weeks after the viral injections to allow stable level of GCaMP6 expression to establish. Under isoflurane anesthesia (3% in room air), the femoral artery and femoral vein were cannulated for the arterial blood pressure recordings and the delivery of drugs, respectively. Then the animals were given α-chloralose to induce and maintain the stable level surgical anesthesia (100 mg kg^−1^ bolus i.v., 30 mg kg^−1^h^−1^ maintenance i.v.). A tracheotomy was performed and the animals were artificially ventilated using a positive pressure rodent ventilator (tidal volume ~8–10 ml kg^−1^; frequency ~60 strokes min^−1^). The body temperature was maintained at 37.0±0.5 °C with a servo-controlled heating blanket. The head of the animal was secured in a stereotaxic frame. Arterial blood samples were taken regularly to monitor blood *P*O_2_, *P*CO_2_ and pH (RAPIDLab 348EX, Siemens). Inspired gas composition and/or rate/volume of the ventilation were adjusted to maintain arterial *P*O_2_ within the range: 100 – 110 mmHg, *P*CO_2_: 35-45 mmHg and pH: 7.35-7.45. To visualize Ca^2+^ responses in NTS astrocytes, the dorsal surface of the brainstem was exposed as described in detail previously (Gourine et al., 2008). [Ca^2+^]_i_ responses in the NTS astrocytes evoked by electrical stimulation of the central end of the vagus nerve (5 s stimulation; 5 Hz, 0.8 mA, 10 ms pulse duration) were recorded using Leica fluorescence microscope and MiCAM02 high-resolution camera (SciMedia). To minimize movement artifacts, the animals were neuromuscularly blocked with gallamine triethiodide (50 mg kg^−1^, i.v.; then 10 mg kg^−1^ h^−1^ i.v.). Since acute changes in blood pressure in response to vagal stimulation were associated with drifts in focal plane affecting image acquisition, the arterial blood pressure was clamped by infusion of a nitric oxide synthase inhibitor Nω-Nitro-L-arginine methyl ester (L-NAME; 10 mg kg^−1^ h^−1^, i.v.) and ganglion blocker chlorisondamine (1 mg kg^−1^, i.v.). Four stimulations were applied: 2 control stimulations followed by stimulations in the presence of increasing doses of 5-HT_2A_ antagonist ketanserin given systemically (100 µg kg^−1^ and 300 µg kg^−1^, i.v.). Stabilisation periods of 10 min between stimulations were allowed. In a separate set of experiments, stimulations were performed in the absence and presence of AMPA receptor antagonist CNQX (10 mM in aCSF; topical application). Imaging data were collected and analysed using MiCaM BV_ANA software.

### Cell culture and calcium imaging in vitro

Primary astroglial cultures were prepared from the cortical, hippocampal, cerebellar, and dorsal brainstem tissue of rat pups (P2–P3 of either sex) as described in detail previously (Kasymov et al., 2013; Angelova et al., 2015; Turovsky et al., 2016). After isolation, the cells were plated on poly-D-lysine-coated coverslips and maintained at 37°C in a humidified atmosphere of 5% CO_2_ and 95% air for a minimum of 12 d before the experiments.

Optical measurements of changes in intracellular calcium ([Ca^2+^]_i_) were performed using an inverted epifluorescence Olympus microscope, equipped with a cooled CCD camera (Retiga; QImaging) as described previously (Kasymov et al., 2013; Angelova et al., 2015; Turovsky et al., 2016).

Experiments were performed in a custom-made flow-through imaging chamber in a standard HBSS balanced with 10 mM HEPES. To visualize [Ca^2+^]_i_ responses, astrocytes were loaded with a conventional Ca^2+^ indicator Fura-2 (5 μM; 30 min incubation; Invitrogen). After incubation with the dye, cultures were washed five times before the experiment. The effects of 5-HT or 5-HT receptor agonists on [Ca^2+^]_i_ in individual astrocytes were recorded. Excitation light was provided by a xenon arc lamp with the beam passing through a monochromator at 340 and 380 nm (Cairn Research) and emitted fluorescence at 515 nm was registered. Imaging data were collected and analysed using Andor software (Andor). All data presented were obtained from at least six separate experiments.

### Recordings of the arterial blood pressure and heart rate using biotelemetry

Systemic arterial blood pressure and heart rate in conscious rats transduced to express dnSNARE or control transgene by the NTS astrocytes were recorded using biotelemetry. The rats were anesthetized with isoflurane (3% in O_2_), a laparotomy was performed, and a catheter connected to a biotelemetry pressure transducer (model TA11PA-C40, DSI) was advanced rostrally into the abdominal aorta and secured in place with Vetbond (3M). The transmitter was secured to the abdominal wall and the incision was closed by successive suturing of the abdominal muscle and skin layers. For postoperative analgesia, the animals received carprofen (4⍰mg⍰kg^−1^ d^−1^; i.p.) for 2 days and were allowed to recover for at least 7 days. After the recovery period and following recordings of the baseline haemodynamic parameters for 24 h, the animals received microinjections of the AVVs to express dnSNARE or control transgene in the NTS astrocytes, as described above. Heart rate and blood pressure were recorded between 7 and 10 days after the injections of the viral vectors when the expression of transgenes reaches the optimal level.

### Analysis of the biotelemetry data

Recordings of the arterial blood pressure were used to calculate heart rate and spontaneous baroreceptor reflex gain (sBRG) for the light and dark periods of the 24h cycle. sBRG was determined from spontaneous changes in systolic blood pressure (SBP) and pulse interval (PI) as described in detail previously (Oosting et al., 1997; Waki et al., 2003). sBRG for reflex bradycardia was calculated over the first 4h of the light phase as the cardiac baroreflex gain in response to spontaneous pressor events has been found to be greater during non-REM sleep compared to awake state (Legramante et al., 2003; Silvani, 2008).

### Assessment of baroreceptor reflex gain (BRG) in anesthetized animals

In anesthetized animals instrumented for the recordings of the arterial blood pressure and heart rate (as described above), baroreceptor reflex was activated by i.v. infusion of norepinephrine (1 µg kg^−1^). Concomitant changes in blood pressure and heart rate were recorded from 3 consecutive stimulations delivered 3 min apart. Baroreflex sensitivity was assessed in the absence and presence of P2Y_1_ receptor antagonist MRS-2500, applied on the dorsal surface of the brainstem (5 µM). The BRG was calculated as a ratio of changes in HR to that of mean arterial blood pressure (bpm/mmHg) for reflex bradycardia, and BRG values were averaged over 3 measurements made in control conditions and after blockade of P_2_Y_1_ receptors in the dorsal brainstem.

### Drugs

5-HT receptor agonists and antagonists were used to identify the type of 5-HT receptors expressed by brainstem astrocytes: 5-HT_2A_ antagonists ketanserin and MDL 100907, 5-HT_2A_ agonist N,N-Dimethyltryptamine (DMT), 5-HT_2B_ agonist BW 723C86, 5-HT_2C_ agonist WAY 161503, 5-HT_3_ antagonist granisetron. Phospholipase C activity was blocked with U73122. P2Y_1_ receptors were blocked with MRS-2500. AMPA receptors were blocked with CNQX. All drugs were purchased from Tocris Bioscience.

### Data analysis

Physiological data obtained in the experiments in anesthetized preparations were recorded and analyzed offline using *Spike2* software (Cambridge Electronic Design). Built-in analysis software tools (Olympus or MiCAM BV_ANA) were used to analyze the results of the imaging experiments. Differences between the experimental groups/treatments were tested for statistical significance by one-way or two-way ANOVA followed by the *post hoc* Tukey–Kramer test or Student’s t test. Data are reported as individual values as well as means ± SEM. Differences with *p*<0.05 were considered to be significant.

## Results

### Vagal afferent simulation activates NTS astrocytes in vivo

Transduction of the NTS under control of the GFAP promoter caused expression of the calcium-sensitive protein GCaMP6 in astrocytes over the mediolateral extent of the NTS (Fig. 1A). Rapid increases in fluorescence intensity (2.0±0.06 ΔF/F_0_; n=5) were observed in response to short (5 s) electrical stimulation of vagus nerve (Fig. 1B). These increases were observed in the area of tissue adjacent to the 4^th^ ventricle, rostral from the calamus scriptorius (Fig. 1C-D, Supplementary Video 1). Corresponding to increases in intracellular [Ca^2+^], the fluorescence changes indicate that astrocytes in the NTS respond to vagal afferent input.

**Figure 1.**
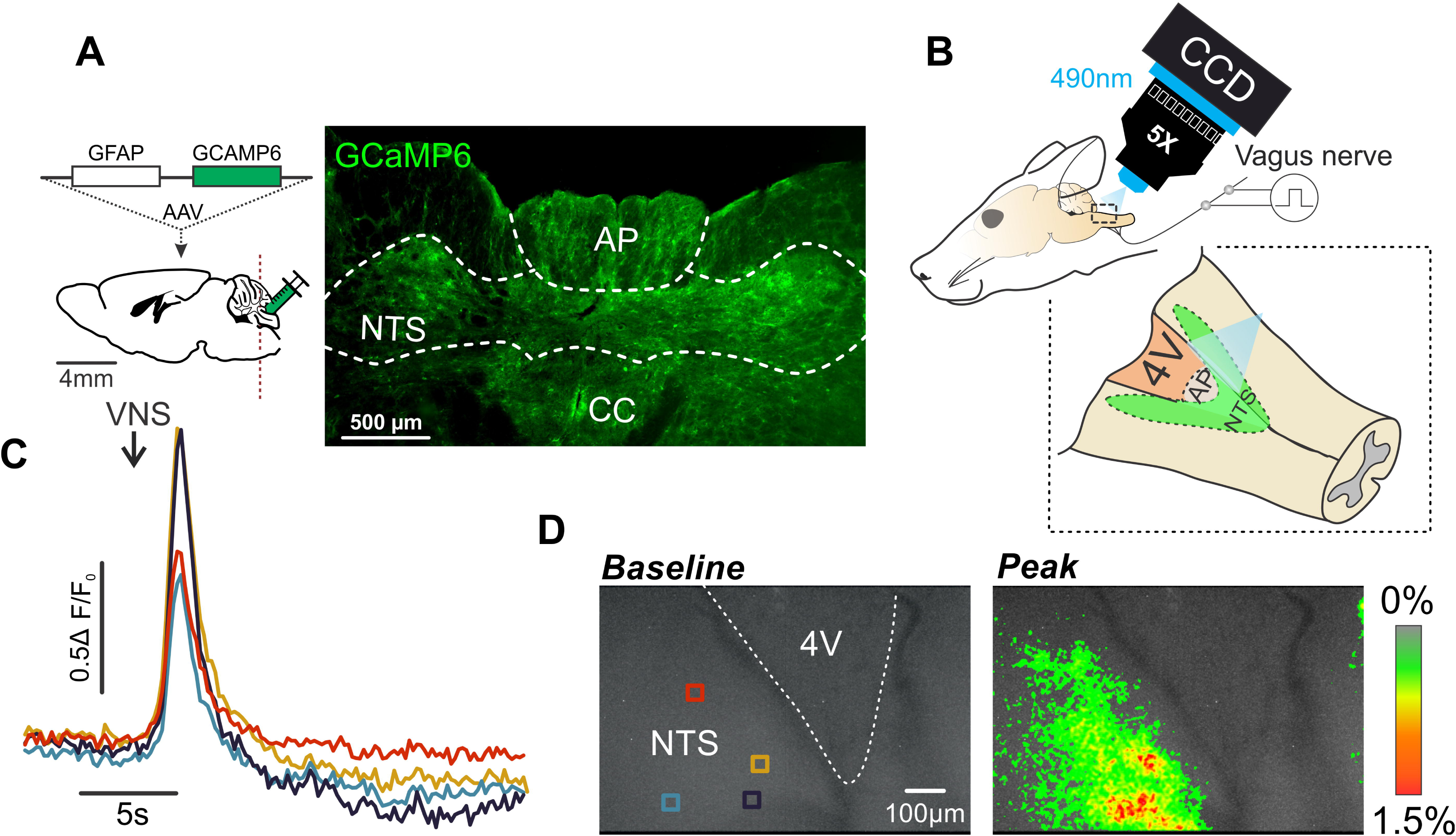
*In vivo* calcium imaging of NTS astrocytes. (***A***) Expression of GCaMP6 (AAV5.GfaABC1D.cytoGCaMP6f.SV40) by NTS astrocytes 4 weeks after transfection. AP, area postrema; cc, central canal. Schematic illustrates the anatomical location of the nucleus of the solitary tract (NTS), vector map and the site of vector delivery. (***B***) Schematic representation of the recording setup; a CCD camera coupled to a fluorescent (excitation wavelength 490nm) microscope records images from the dorsal aspect of the brainstem at the caudal extent of the 4^th^ ventricle (4V); the vagus nerve, mounted on a sliver hook electrode, is stimulated using a constant current source. (***C***) Representative changes in GCaMP6 fluorescence in 4 areas of interest on the dorsal aspect of the brainstem during a 5 s electrical vagus nerve stimulation (VNS). (***D***) False color image of Δ*F/F*_0_ peak intensity changes before vagus stimulation (baseline) and at the peak of fluorescent change during stimulation. Colored boxes indicate regions of interest in (C).

### Brainstem astrocytes express functional 5-HT_2A_ receptors

As 5-HT is known to be co-released with glutamate from vagal afferents, we tested for the presence of functional 5-HT receptors in cultured brainstem astrocytes. These cells responded to application of 5-HT (10 μM) with profound elevations in intracellular [Ca^2+^] (0.164±0.022 fura-2 ratio above baseline, n=10; Fig. 2A). 5-HT-induced Ca^2+^ responses were not affected in the absence of extracellular Ca^2+^ (Ca^2+^-free medium with the addition of 0.5 mM EGTA) (0.115±0.022, n=10, *t*-test, p=0.09; Fig. 2A), suggesting that 5-HT actions recruit Ca^2+^ from the intracellular stores, likely to be mediated by G_q_-coupled 5-HT_2_ receptor subtype (Hoyer et al., 2002). Indeed, [Ca^2+^]_i_ responses triggered by 5-HT in brainstem astrocytes were abolished in the presence of phospholipase-C (PLC) inhibitor U73122 (5 μM; 0.011±0.002 vs 0.340±0.054, n=15, *t*-test, p<0.001; Fig. 2B) and 5-HT_2A_ receptor antagonist ketanserin (0.010 μM; 0.015±0.004 vs 0.172±0.014, n=16, t-test, p<0.001) (Fig. 2C). Neither 5-HT_2B_ agonist BW723C86 (in concentrations 0.001-1 μM) nor 5-HT_2C_ agonist WAY161503 (in concentrations 0.01-5 μM) had an effect on [Ca^2+^]_i_ in brainstem astrocytes (Fig. 2D, Fig. 2E). These data indicated that responses of brainstem astrocytes to 5-HT are mediated by 5-HT_2A_ receptors.

**Figure 2.**
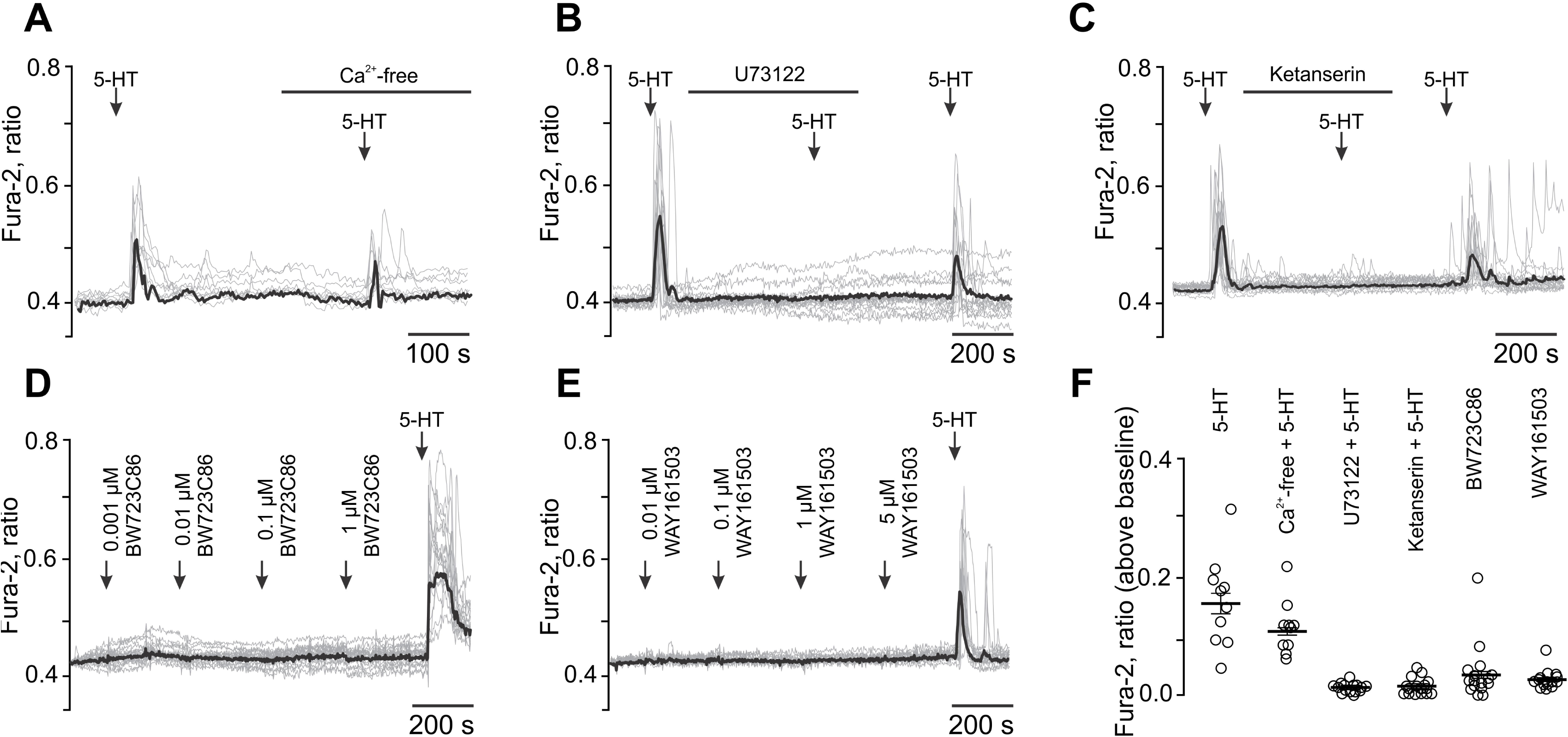
Brainstem astrocytes express functional 5-HT_2A_ receptors. (***A***) Astrocytes cultured from brainstem responded to 5-HT (10 μM) with vigorous elevations in intracellular [Ca^2+^]_i_; 5-HT induced [Ca^2+^]_i_ responses persist in Ca^2+^-free medium. (***B***) 5-HT induced [Ca^2+^]_i_ responses are blocked by PLC-inhibitor U73122 and (***C***) by 5-HT_2A_ antagonist ketanserin. (***D***) Brainstem astrocytes do not respond to 5-HT_2B_ agonist BW723C86 and (***E***) 5-HT_2c_ receptor agonists WAY161-503. (***F***) Summary of maximum [Ca^2+^]_i_ responses to 5-HT and 5-HT receptor agonists in brainstem astrocytes.

Analysis of Ca^2+^ responses induced by 5-HT in astrocytes residing in other areas of the brain (cerebellum, hippocampus and cortex) suggested that the profile of 5-HT receptors expressed by brainstem astrocytes is distinct from that of the forebrain astrocytes. Similar to brainstem astrocytes, cerebellar astrocytes (Bergmann glia) express 5-HT_2A_ receptors, while Ca^2+^ responses induced by 5-HT in cortical and hippocampal astrocytes are mediated by 5-HT_3_ receptors and Ca^2+^ entry from the extracellular space (see Fig. S1 for detailed pharmacological analysis).

### 5-HT_2A_ receptors mediate Ca^2+^ responses of NTS astrocytes evoked by vagal afferent simulation

To determine if the identified glial 5-HT_2A_ receptors are functional *in vivo*, calcium transients in response to vagus nerve stimulation were recorded in NTS astrocytes expressing calcium-sensitive protein GCaMP6. The 5-HT_2A_ receptor antagonist ketanserin markedly decreased the amplitudes of [Ca^2+^]_i_ responses evoked by stimulation in a dose dependent manner (ΔF/F_0_=1.3±0.03 and 1.0±0.02 following administration in doses of 100 µg kg^−1^ and 300 µg kg^−1^, respectively; n=5; p<0.05; Fig. 3A-B, Supplementary Video 2). However, when administered topically, the AMPA receptor antagonist CNQX completely abolished calcium responses of NTS astrocytes evoked by vagus nerve simulation (Fig. 3D).

**Figure 3.**
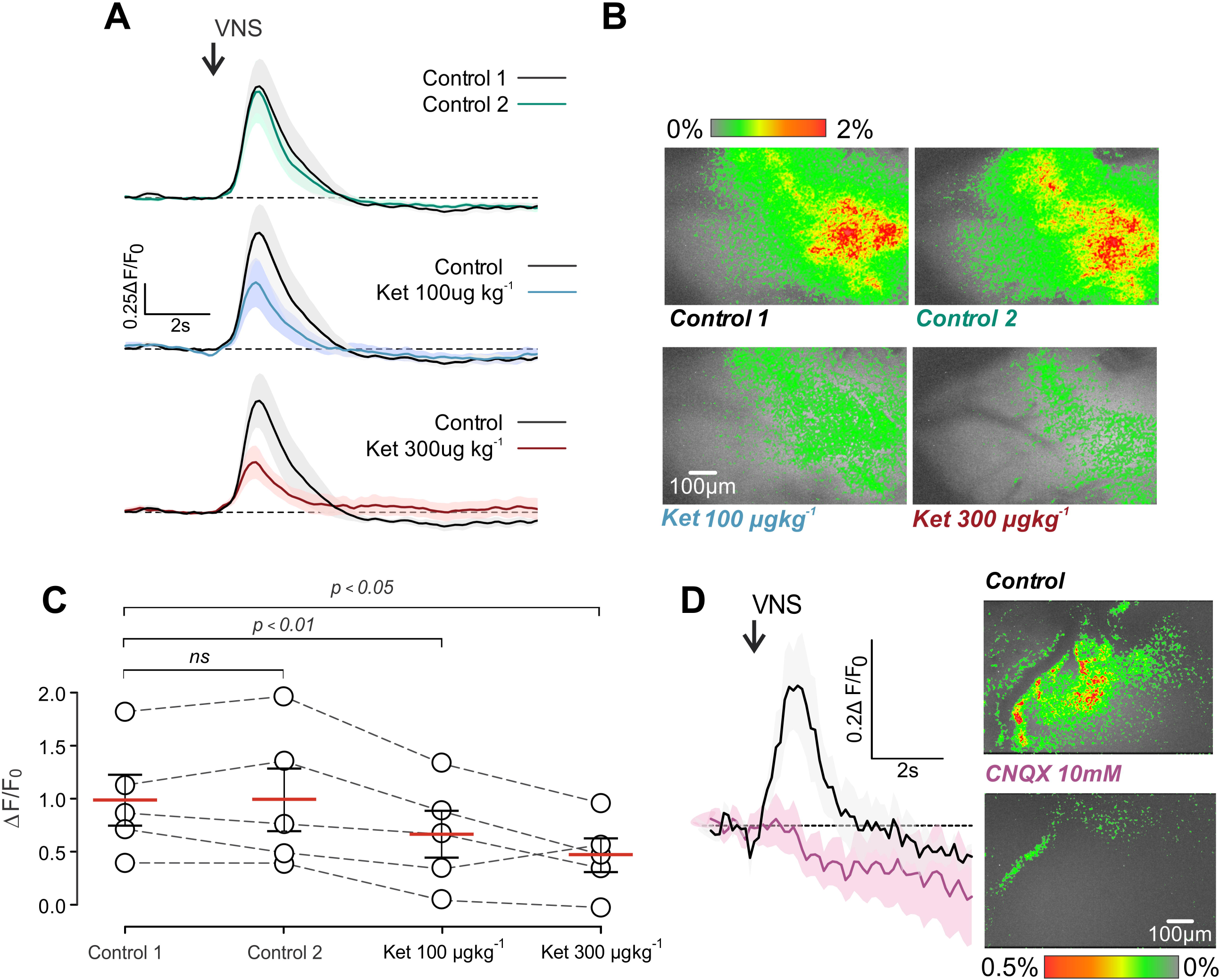
NTS astrocytes respond to 5-HT and glutamate released by vagal afferent stimulation *in vivo*. (***A***) GCaMP6 fluorescence changes (±S.E.M) showing [Ca^2+^]_i_ dynamics in response to vagal nerve stimulation in control conditions separated by 10 min intervals and in presence of the increasing doses of ketanserin (100-300μg kg^−1^; n=5) (***B***) Representative false color images of peak fluorescence intensity changes during control conditions and increasing doses of ketanserin. (***C***) Grouped data showing average (±SEM) peak Ca^2+^-related fluorescence intensity changes in the NTS astrocytes that are inhibited by 5-HT_2A_ antagonist ketanserin: intravenous administration of ketanserin significantly (p<0.01) decreased the amplitude of [Ca^2+^]_i_ responses in a dose-dependent manner (n=5) when compared with repeated measures (one-way ANOVA followed by Holm-Sidak’s multiple comparisons test). (***D***) Average astroglial [Ca^2+^]_i_ responses induced by as determined by changes in (±SEM) before and after topical application of AMPA receptor antagonist (10mM; n=4). Insert shows a representative false color image of Δ*F/F*_0_ peak intensity changes during vagal stimulation before and after CNQX application.

### Blockade of vesicular release mechanisms in NTS astrocytes alters baroreflex sensitivity

To determine the functional significance of NTS astroglial Ca^2+^ responses induced by afferent stimulation, we next determined whether blockade of Ca^2+^-dependent vesicular release mechanisms in these cells have an effect on baroreflex control. In conscious freely moving rats, the expression of dnSNARE in the NTS astrocytes (Fig. 4A) was associated with a significant reduction in baroreflex sensitivity 7 and 10 days after the injections of the viral vectors, when the expression of the transgene peaked (Fig. 4B, sBRG 1.7±0.11 and 1.5±0.10 bpm/mmHg vs 1.0±0.10 bpm/mmHg, p<0.001). Baroreflex sensitivity was unaffected in animals transduced to express the control transgene in the NTS astrocytes (Fig. 4B, sBRG 1.1±0.08 and 1.1±0.13 bpm/mmHg vs 1.0±0.07 bpm/mmHg, p>0.05).

**Figure 4.**
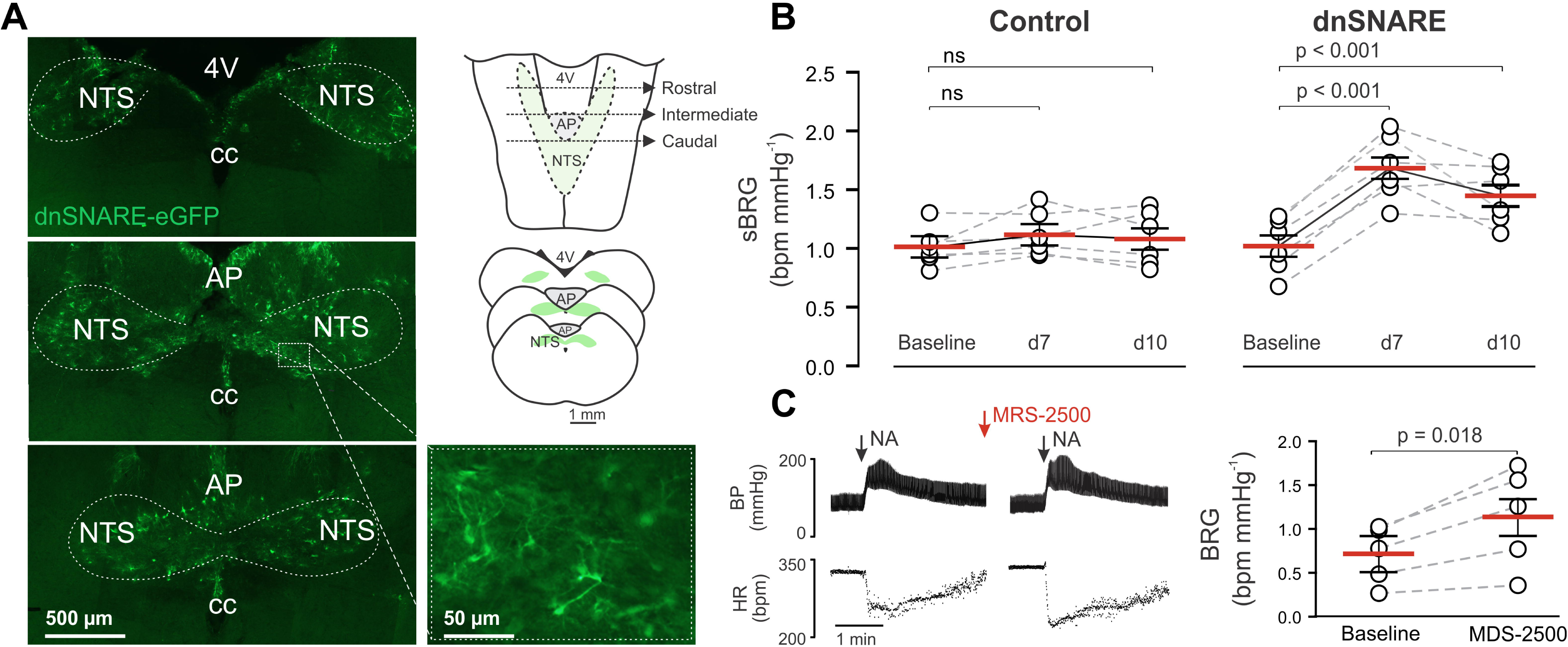
Blockade of vesicular release or P2Y_1_ receptors in NTS astrocytes increases baroreflex sensitivity. (***A***) Expression of dnSNARE-eGFP by astrocytes in rostral, intermediate and caudal NTS 10 days after transfection (left) and schematic drawings of the rat brainstem illustrating position of the sections (right). Higher-magnification image of dnSNARE-eGFP expression (green) in astrocytes of the intermediate NTS (bottom right). AP, area postrema; cc, central canal (***B***) In animals transduced to express dnSNARE in the NTS astrocytes, the sBRG significantly increased when assessed on days 7 and 10 after viral injections compared to baseline values. (***C***) In anesthetized animals, blockade of NTS P2Y_1_ receptors with topical application of 5 μM MRS-2500 augmented bradycardia induced by baroreceptor activation with systemic norepinephrine (NA).

### Inhibition of baroreflex sensitivity at the level of NTS is mediated by P2Y_1_ receptors

Increases in [Ca^2+^]_i_ lead to Ca^2+^-dependent vesicular release of ATP by astrocytes (Gourine and Kasparov, 2011). We hypothesised that NTS inhibitory interneurons could be activated by ATP released by astrocytes via activation of P2Y_1_ receptors, as a similar P2Y receptor-mediated inhibitory mechanism has been shown to modulate neuronal excitability in the cerebral cortex (Wang et al., 2012). To test the involvement of the P2Y_1_ receptor, baroreflex sensitivity was assessed in anesthetized animals before and after application of a potent and selective P2Y_1_ receptor antagonist MRS-2500 topically to the brainstem (5 µM; (Kim et al., 2003). The baroreflex was activated by bolus injections of norepinephrine (NA, 1 µg kg^−1^). It was found that blockade of NTS P2Y_1_ receptors significantly increased the baroreflex gain (Fig. 4C, 1.1±0.26 bpm/mmHg vs 0.7±0.15 at baseline, p=0.018).

## Discussion

The importance of astrocytes in the operation of the NTS circuits has been suggested previously by (Lin et al., 2013), who reported that ablation of astrocytes using toxin saporin results in cardiovascular reflex attenuation, lability of arterial pressure, damage of cardiac myocytes and, in some animals, sudden cardiac death. However, considering the important role played by astrocytes in providing structural and metabolic support, as well as K^+^ buffering and glutamate re-cycling, it is not surprising that ablation of astrocytes has a major impact on the neuronal function. Therefore, the role of astrocytes in the subtleties of neuronal information processing and afferent integration within the NTS remain unknown. In this study we aimed to determine the role of astrocytes in the NTS mechanisms that process cardiovascular afferent information.

First, *in vivo* calcium imaging demonstrated that NTS astrocytes respond to vagal afferent input with increases in intracellular [Ca^2+^]. This is due, in part, to glutamate release from vagal afferent terminals and is in agreement with the *in vitro* observations by McDougal and co-workers (2011), who reported that NTS astrocytes respond with increases in [Ca^2+^]_i_ to stimulation of the solitary tract (McDougal et al., 2011). As 5-HT is a known co-transmitter released from vagal afferent terminals, we reasoned that it could partially shape this response. Accordingly, it was demonstrated that cultured brainstem astrocytes respond to 5-HT with robust [Ca^2+^]_i_ increases which were blocked by 5-HT_2A_ antagonist ketanserin (Bonhaus et al., 1995) or by inhibition of phospholipase C activity (Fig.2). Although 5-HT_3_ receptors have been previously reported to be expressed by NTS astrocytes (Huang et al., 2004), we found no evidence of their involvement in mediating the actions of 5-HT. NTS astrocytes appear to be distinct from the forebrain astrocytes (cortical and hippocampal) where 5-HT effects are mediated solely by ionotropic 5-HT_3_ receptor (Fig. S1). These data suggested that the brainstem astrocytes can detect the release of 5-HT by the afferent projections terminating in the NTS.

We confirmed that 5-HT_2A_ receptors were of functional significance when modulating intracellular calcium *in viv*o: peak [Ca^2+^] responses in NTS astrocytes are reduced by ∼50% in the presence of 5-HT_2A_ antagonist ketanserin (Fig 3). 5-HT released from vagal afferents alone was unable to cause a detectable increase in astrocyte [Ca^2+^]_i_, as blockade of NTS AMPA receptors completely abolished this effect of vagal nerve stimulation on astrocytes. It is also possible that 5-HT is released from other sources within the NTS, as suggested previously (Hosford et al., 2015), not exclusively from vagal afferents. Tyrosine hydroxylase-expressing cells of the brainstem raphe nuclei have reciprocal connections with the NTS and could be activated by projections from the NTS (Thor and Helke, 1989; Rosin et al., 2006).

Finally, the key experiment involved specific targeting of the Ca^2+^-dependent vesicular release machinery in the NTS astrocytes to determine the functional significance of astroglial signaling in operation of a key homeostatic reflex – the baroreflex (Fig. 4). It was found that inhibition of this specific astroglial signaling pathway significantly increased the baroreflex sensitivity in awake behaving rats. The main astroglial signaling molecule ATP is known to inhibit local neuronal circuits indirectly following breakdown to adenosine, – the mechanism first reported to operate in retina (Newman, 2003). Indeed, activation of adenosine A_1_ receptors in the NTS inhibits the baroreceptor depressor reflex (Scislo and O’Leary, 2005). However, the data obtained in this study suggested the existence of a different mechanism. It was found that pharmacological blockade of metabotropic P2Y_1_ receptors in the NTS has a similar effect on baroreceptor reflex control as dnSNARE-mediated inhibition of Ca^2+^-dependent vesicular release in astrocytes. This suggests that, upon activation, NTS astrocytes release ATP which is acting on NTS inhibitory neurons expressing P2Y_1_ receptors, – the mechanism analogous to that described by (Wang et al., 2012) in the cortex.

Baroreceptor reflex is critically important for the short term (or beat-to-beat) control of the arterial blood pressure. There is strong evidence that impaired baroreflex function contributes to the development of cardiovascular disease and serves as a robust predictor of cardiovascular and all-cause mortality (La Rovere et al., 1998; La Rovere et al., 2001; McCrory et al., 2016). The mechanisms underlying impairment of baroreflex function in pathological conditions may result from several peripheral and central alterations, many of which are still unknown. Among central mechanisms, the activation of the cardiac sympathetic afferent reflex which alters the baroreflex via angiotensin II (ANG II) type 1 (AT1) receptors in the NTS (Gao et al., 2005; Wang et al., 2007), and reduction of brain-derived neurotropic factor (BDNF) neurotransmission in the NTS (in particular, reduced BDNF-tyrosine receptor kinase B signaling) (Becker et al., 2016) were proposed. The results of the present study offer another plausible mechanism. The many different pathological conditions that are associated with the development of the systemic inflammatory response leading to the activation of NTS astroglia would be expected to facilitate the release of ATP, increase the basal concentration of ATP in the extracellular milieu and inhibit the baroreflex centrally. Activation of astrocytes and reactive astrogliosis have been reported under conditions of CNS trauma, infection, ischaemia, stroke, autoimmune responses, and neurodegenerative disease (e.g. Alzheimer’s, Parkinson’s disease) (reviewed by Sofroniew and Vinters, 2010). This would be expected to reduce the baroreflex sensitivity via P2Y_1_ receptor-mediated NTS mechanism described here. Additionally, repeated activation of chemosensory inputs has been shown to chronically inhibit the baroreflex and is thought to contribute to the pathology of conditions such as sleep apnoea (see Mifflin et al., 2015). Activation of chemosensory inputs increases extracellular 5-HT concentration within the NTS via vagal/hypoglossal afferent release, and also by inputs from central chemosensory nuclei (Kellett et al., 2005; Wu et al., 2019). This would have the effect of amplifying the activation of astrocytes and further decreasing baroreflex sensitivity.

In conclusion, the data obtained in the present study suggest that astrocytes are integral components of the NTS mechanisms which process incoming afferent information. They are activated by glutamate (McDougal et al., 2011) and 5-HT, also released by vagal afferent fibres and acting on 5-HT_2A_ receptors (Fig. 5). Astrocyte activation appears to be functionally significant for the operation of the aortic baroreceptor reflex. These data suggest that activation of astrocytes in response to afferent stimulation leads to the vesicular release of ATP acting on P2Y_1_ receptors on local interneurons to modulate the baroreflex sensitivity. This study adds to the growing body of evidence supporting an active role of astrocytes in the information processing in the central nervous system.

**Figure 5.**
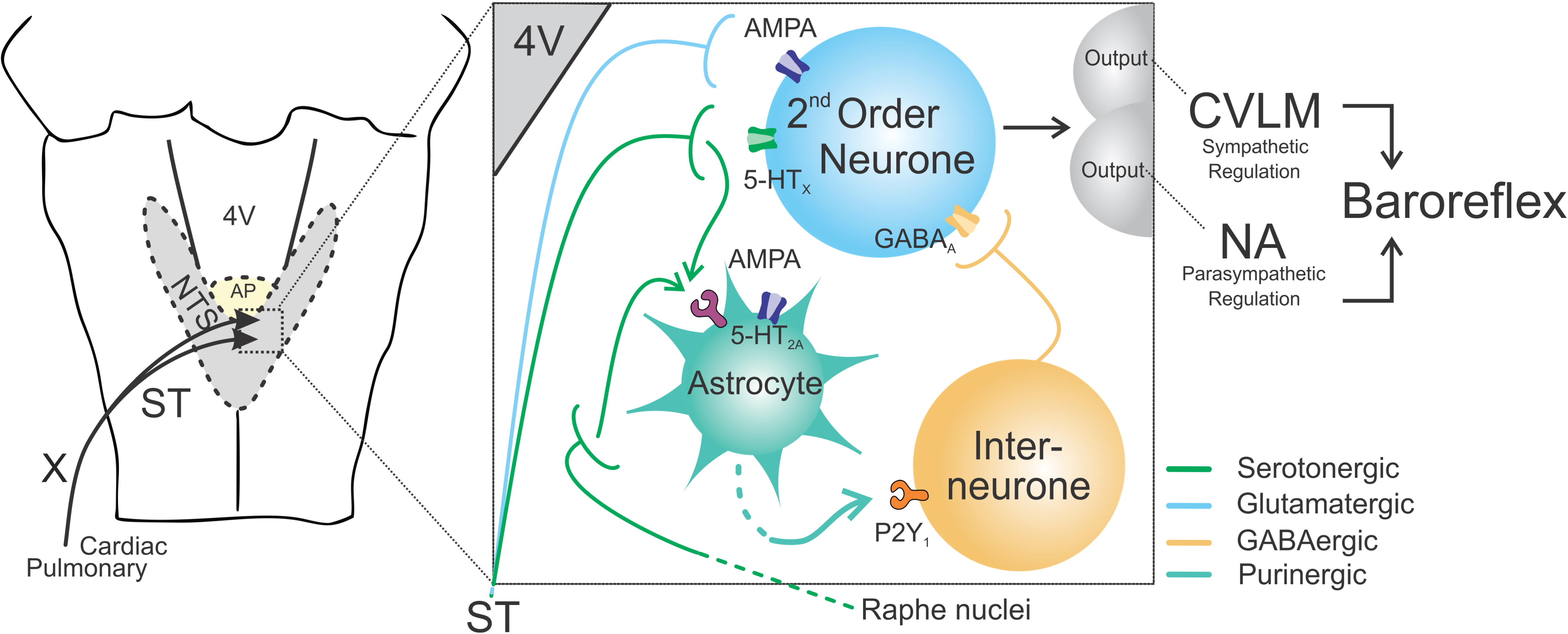
Schematic representation of the proposed circuit involved in the control of baroreflex at the level of NTS. Vagal afferent neurons release 5-HT and glutamate acting on both 2^nd^ order neurons and astrocytes in the NTS. 5-HT acting at 5-HT_2A_ receptors mediates (at least in part) the effects of vagal afferent activity on NTS astrocytes. In response to incoming afferent information, NTS astrocytes release ATP which restricts the expression of baroreflex via activation of P2Y_1_ receptors on local inhibitory interneurons.

Supported by British Heart Foundation (Refs: PG/13/79/30429 and RG/14/4/30736), The Wellcome Trust (to AVG; Refs: 200893) and Marie Curie Fellowship (to SM; Ref: 654691).

---

***Supplementary Figure 1.*** Distinct 5-HT receptors mediate serotonin-induced [Ca^2+^]_i_ responses in astrocytes residing in different parts of the central nervous system. (***A***) **Cerebellar astrocytes** express 5-HT_2A_ receptor. (***Ai***) 5-HT induced [Ca^2+^]_i_ responses in cerebellar astrocytes are blocked by PLC-inhibitor U73122 which suggests expression of 5-HT_2_ receptor subtype on cerebellar astrocytes. (***Aii***) [Ca^2+^]_i_ responses in cerebellar astrocytes are blocked by 5-HT_2A_/5-HT_2C_ antagonist ketanserin. (***Aiii***) cerebellar astrocytes do not respond to 5-HT_2c_ receptor agonist WAY161503. (***B***) **Hippocampal astrocytes** express 5-HT_3_ receptor. (***Bi***) 5-HT evoked [Ca^2+^]_i_ responses in hippocampal astrocytes persist in the presence of PLC-inhibitor U73122 which suggests expression of 5-HT_3_ receptor subtype, which is ligand-gated ion channel and the only 5-HT receptor not associated with PLC. (***Bii***) 5-HT evoked [Ca^2+^]_i_ responses in hippocampal astrocytes persist in the presence of 5-HT_2A_/5-HT_2C_ antagonist ketanserin. (***Biii***) All 5-HT evoked [Ca^2+^]_i_ responses in hippocampal astrocytes are blocked by 5-HT_3_ antagonist granisetron. (***C***) **Cortical astrocytes** express 5-HT_3_ receptor. (***Ci***) 5-HT evoked [Ca^2+^]_i_ responses in cortical astrocytes persist in the presence of PLC-inhibitor U73122. (***Cii***) Cortical astrocytes do not respond to 5-HT_2_ receptor agonist N,N-Dimethyltryptamine (DMT). (***Ciii***) All 5-HT evoked [Ca^2+^]_i_ responses in cortical astrocytes are blocked by 5-HT_3_ antagonist granisetron.

***Supplementary Video 1.*** Representative recording of astrocytic intracellular calcium activity in the NTS during vagal nerve stimulation under control conditions.

***Supplementary Video 2.*** Representative recording of astrocytic intracellular calcium activity in the NTS during vagal nerve stimulation 10 min after application of 300μg kg^−1^ ketanserin (i.v).

