## Supplementary figures and images for "Astrocytes modulate baroreceptor reflex sensitivity at the level of the nucleus of the solitary tract"

### Supplementary Figure 1

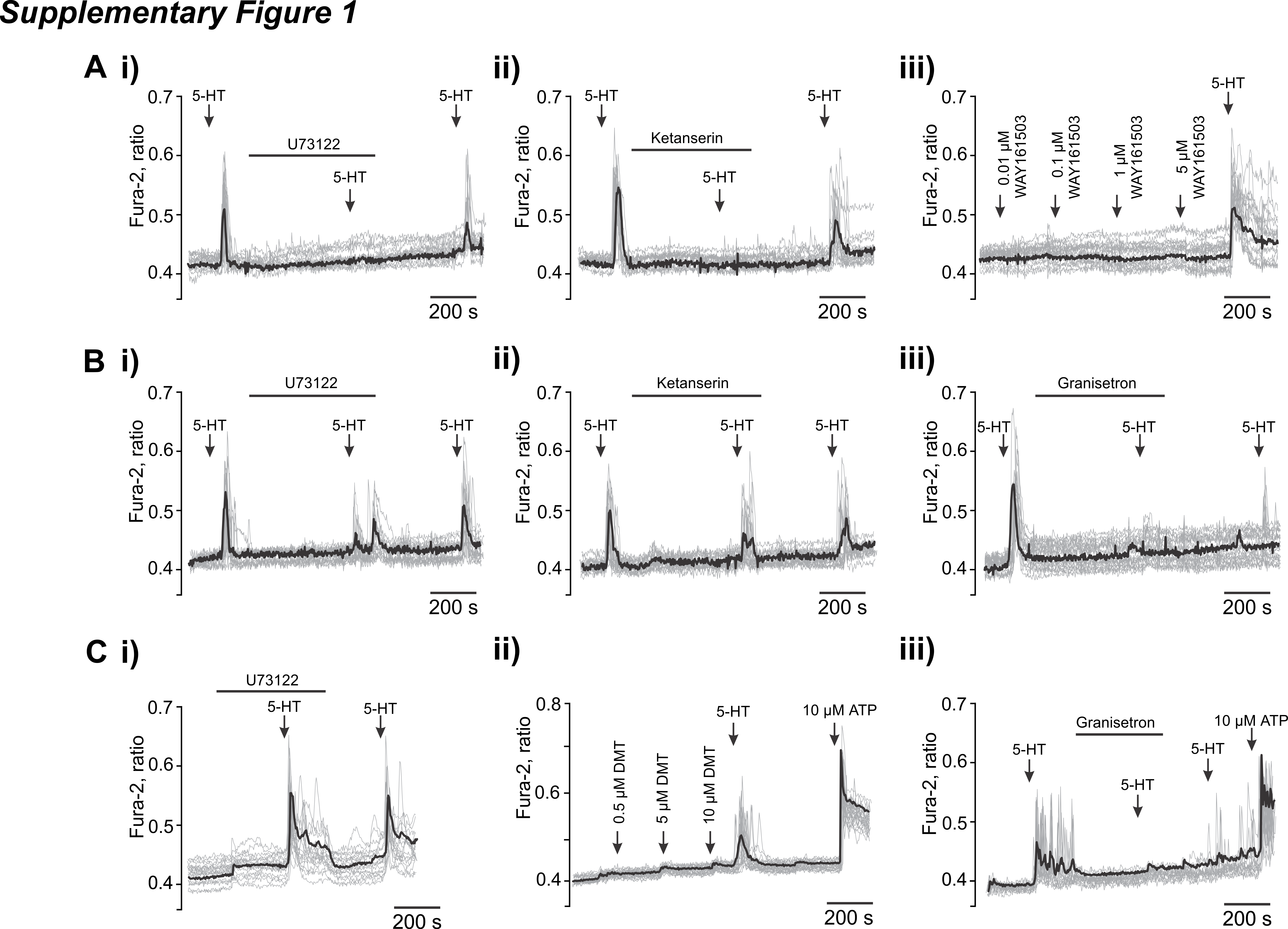
